## supplemental figures for "A standardized gnotobiotic mouse model harboring a minimal 15-member mouse gut microbiota recapitulates SOPF/SPF phenotypes"

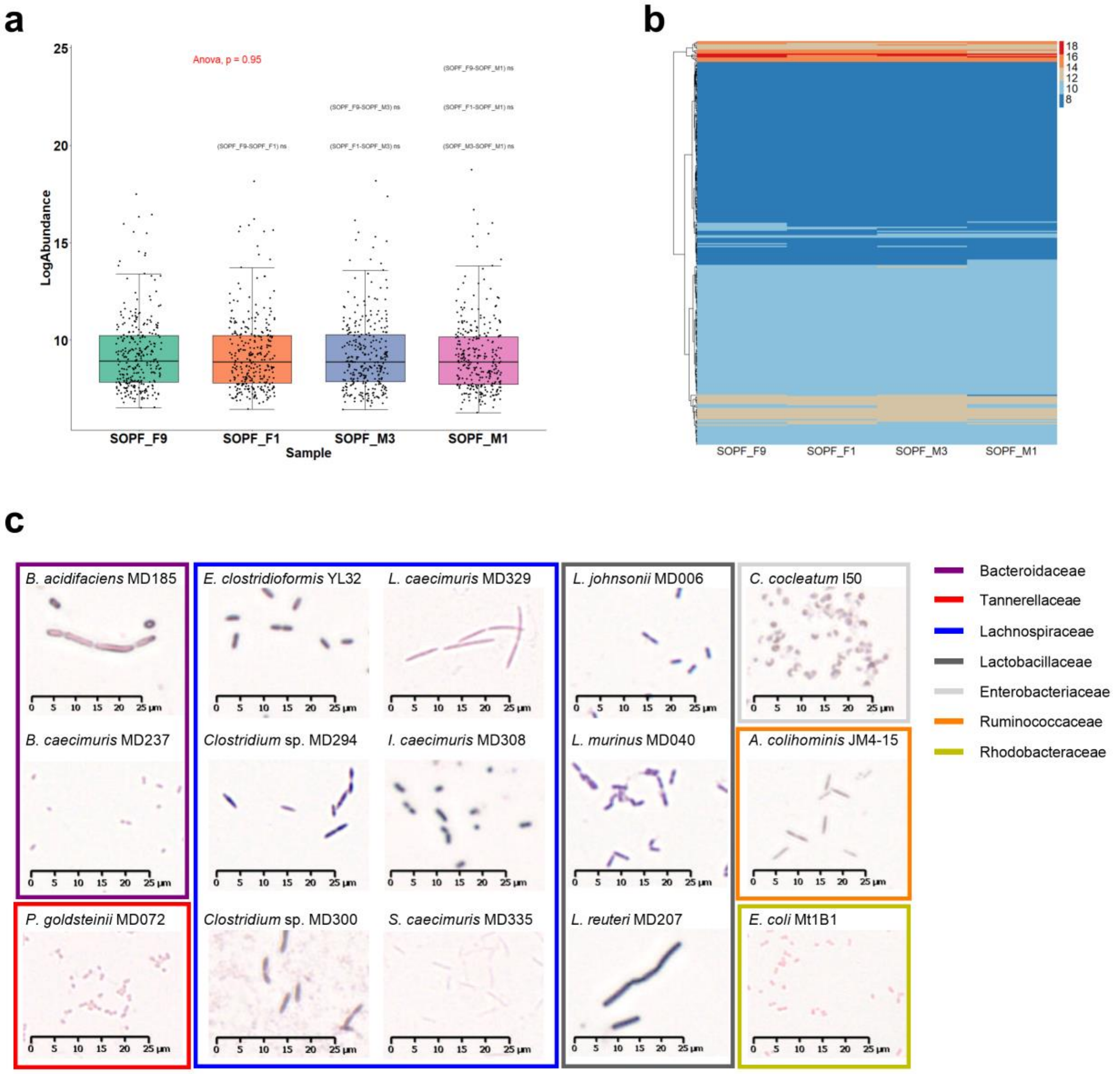


**Additional file 1: Fig. S1. Distribution of bacterial families among the C57BL/6J SOPF mice used to design the GM15 community and morphological features of the GM15 strains. a** Box plots (median, and 25$^{th}$ and 75$^{th}$ percentiles) with whiskers (lower and upper value within the 1.5 fold of the interquartile range) where dots represent individual bacterial families. ANOVA and t test. **b** Heatmap representing the distribution and classification of the log-normalized abundance of each bacterial family (lines) for each mouse (columns). **c** Each bacterial strain was grown individually from a single colony isolated on agar medium and amplified in liquid culture to exponential growth phase. Bacteria were Gram-stained and imaged by light microscopy (80-fold magnification, NanoZoomer S60, Hamamatsu). Members of *Bacteroidaceae*, *Tannerellaceae* and *Enterobacteriaceae* stained Gram-negative, while

*Lactobacillaceae*, *Erysipelotrichaceae* and *Ruminococcaceae* stained Gram-positive. *Lachnospiraceae* stained Gram-positive, except *Longibacillum caecimuris* MD329 and *Subtilibacillum caecimuris* MD335, which stained Gram-negative likely due to their cell wall structure as already reported for other Clostridiales [95].

**Additional file 2: Table S1. KEGG clusters.**

**Additional file 3: Table S2. GM15 strains-specific primers.** Sequences of primers designed in this study and detection limits by qPCR microfluidic assay.

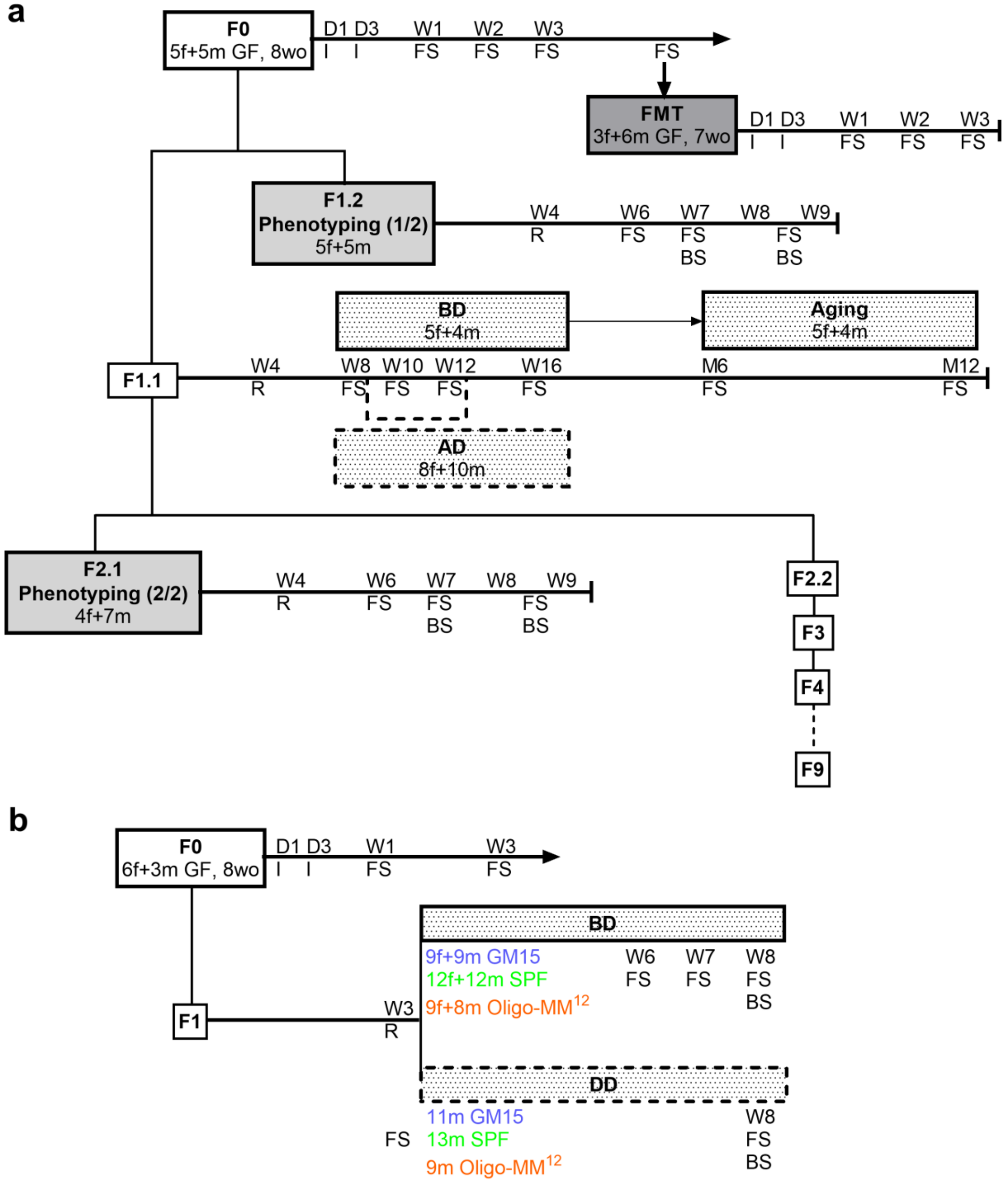


AD: alternative diet; BD: breeding diet; BS: blood sampling; D: day; DD: depleted diet; f: female; F: filial generation; FMT: fecal microbiota transplantation; FS: fecal sampling; GF: germ-free; I: inoculation of bacteria; m: male; M: month; R: randomisation at weaning; SPF: specific pathogen-free; W: week; wo: weeks old

**Additional file 4: Fig. S2. *In vivo* experimental design. a** Facility 1. First, 5 couples of 8 week-old C57BL/6J GF mice were colonized by oral gavage with the fresh frozen GM15 bacterial community, twice at 48h interval (F0). GM15 microbiota stability was assessed by qPCR microfluidic assay from feces collected after 1, 2 and 3 weeks. Then, in order to evaluate the reproducible transfer of the GM15 microbiota by fecal microbiota transplantation, 7 week-old C57BL/6J GF mice were orally gavaged with a suspension of fresh fecal pellets from GM15 mice (F0), twice at 48h interval. Again, qPCR microfluidic assay from feces collected after 1, 2 and 3 weeks was carried out. Next, the GM15 mouse line was amplified to monitor the GM15 microbiota through nine filial generations at 6 weeks of age (F1-F9), and allow the phenotyping study from two consecutive generations (F1.2 and F2.1). Reproduction performance and perinatal mortality were recorded, 4 week-old mice were randomly selected at weaning, monitored weekly for body weight and size, and feed intake, until sacrifice at 8-9 weeks of age. GF and SOPF mice were also studied as control groups. Besides, a comparative analysis was done on the fecal microbiota of 8 week-old GM15 mice (F1.1) either fed with the breeding diet or an alternative isocaloric diet given for 4 weeks. Finally, the fecal microbiota of 6 month-old and 12 month-old control GM15 mice fed with the breeding diet was analyzed. **b** Facility 2. First, 3 trios of 8 week-old C57BL/6J GF mice were colonized by oral gavage with the fresh frozen GM15 bacterial community, twice at 48h interval (F0). GM15 microbiota stability was assessed by qPCR microfluidic assay from feces collected after 1 and 3 weeks. Next, the GM15 mouse colony was amplified to monitor the GM15 microbiota in generation F1 at 6 weeks of age, allow the phenotyping study and assess mice' response to diet-induced stunting.

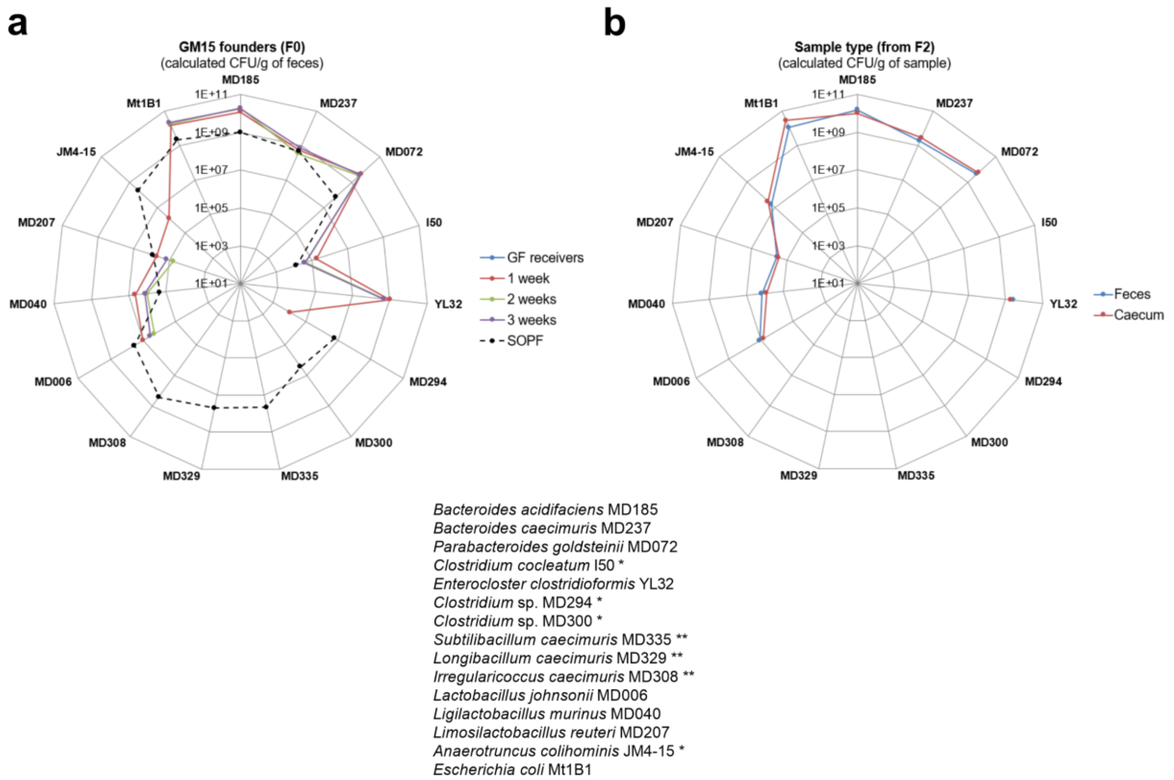


**Additional file 5: Fig. S3. Assessment of gut microbiota stability in GM15 founders (F0, n=10), and of reproducibility between fecal and caecal samples of individual GM15 mice (F2, n=11).** SOPF group shows the distribution of each GM15 strains in the complex gut microbiota of 8-week-old SOPF mice. The absolute quantification of each strain was determined by specific qPCR microfluidic assay. * Strains I50, MD294, MD300 and JM4-15 were at the detection limit of the qPCR microfluidic assay, and thus were not detected in all samples. ** Strains MD335, MD329 and MD308 were below detection limit of the qPCR microfluidic assay. Strain YL32, obtained from the DSMZ collection, was not detected in our SOPF colony. **a** Radar plot showing the GM15 strains distribution in feces of C57BL/6J GF mice before and 1, 2 and 3 weeks after the oral colonization with the GM15 community. **b** Radar plot showing the reproducible detection of the GM15 strains in feces and caecum collected from the same GM15 mice.

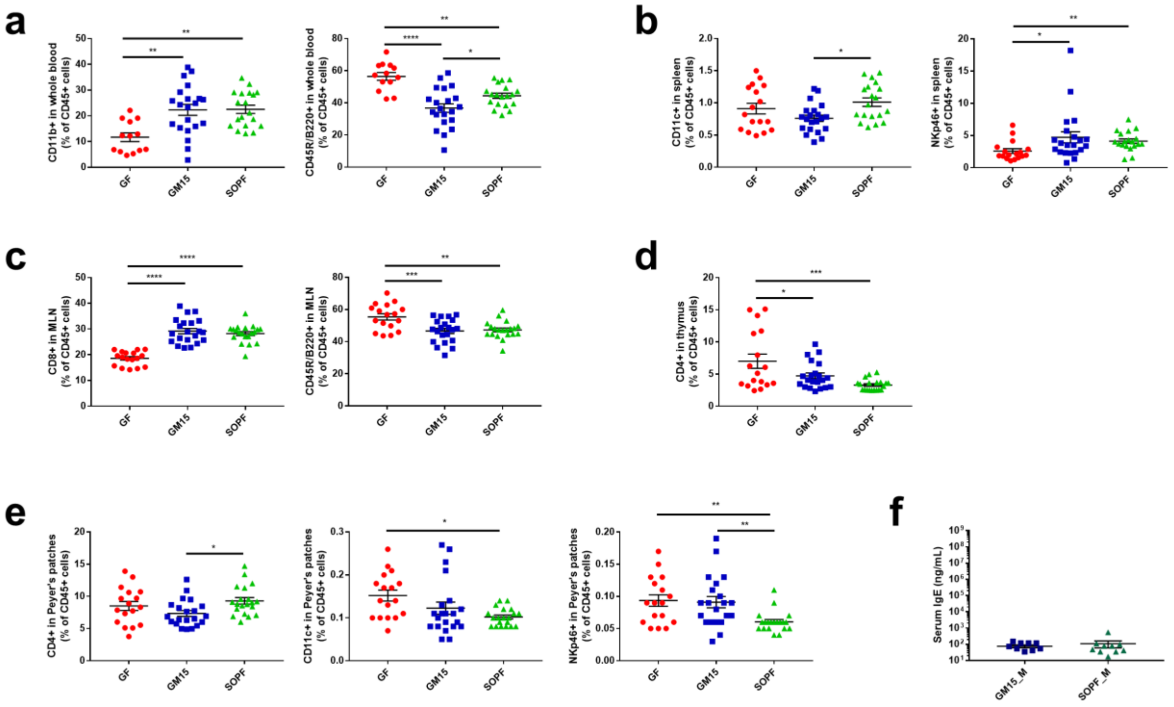


**Additional file 6: Fig. S4. Immune cell populations profiling analysis by flow cytometry in different organs.** Data are represented as % of CD45+ cells. **a-e** Dot plots where dots, lines and error bars represent respectively individual mice (13-17 GF, 21 GM15 and 19-20 SOPF), means and SEM. **a** Monocytes and B cells population in whole blood. Tukey's multiple comparison analyses. **b** DC and HN cells in spleen. Tukey's multiple comparison analysis (DC) and Dunn's multiple comparison analysis (NK cells). **c** CD8+ T cells and B cells in MLN. Tukey's multiple comparison analyses. **d** CD4+ T cells in thymus. Tukey's multiple comparison analysis. **e** CD4+ T cells, DC and NK cells in PP. Tukey's multiple comparison analysis (CD4+ T cells) and Dunn's multiple comparison analyses (DC and NK cells). **f** IgE ELISA assay (10 GM15_M and 10 SOPF_M). Mann-Whitney test. *P<0.05, **P<0.01, ***P<0.001, ****P<0.0001.

**Additional file 7: Table S3. Polar metabolites and lipids concentration in plasma samples.** The metabolites quantification was performed with the help of Chenomx NMR suite 8.6. The non-polar database profiles were created with the help of Compound builder module using $^{1}$H NMR spectra of authentic lipid standards.

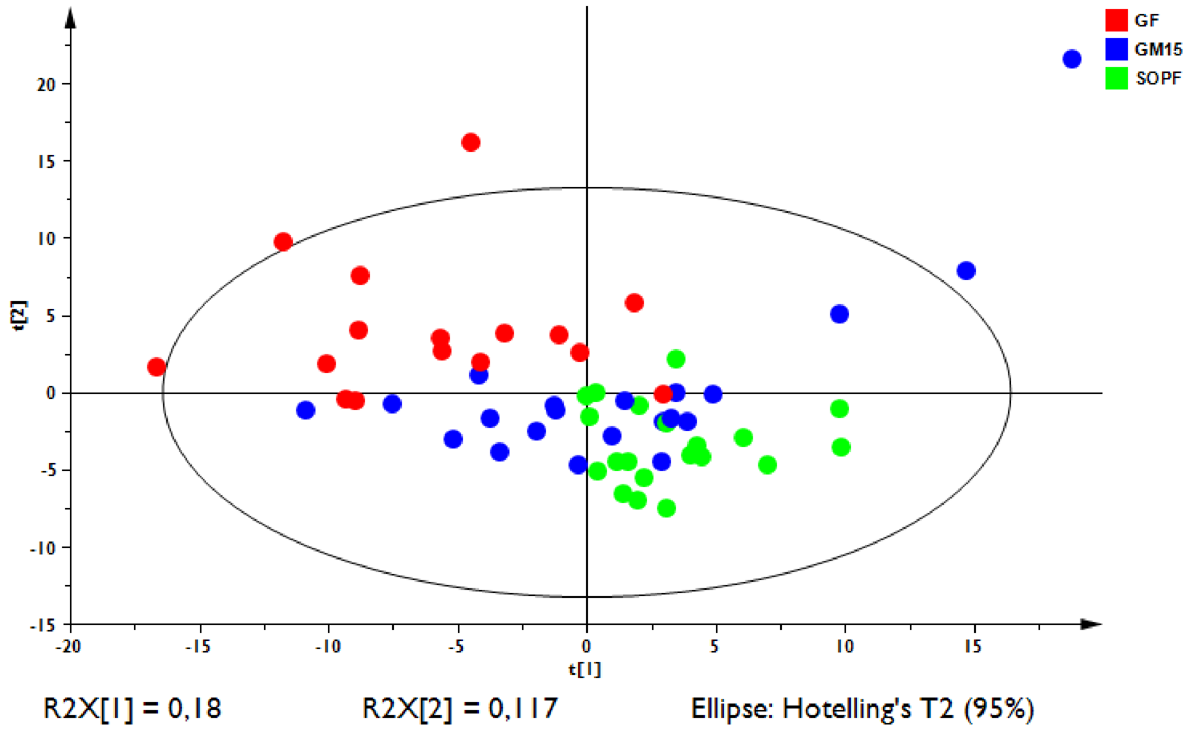


**Additional file 8: Fig. S5. PCA screen plot based on $^{1}$H NMR plasma fingerprints.** Processed $^{1}$H NMR plasma spectra of polar metabolites were binned with AMIX v3.9.14 software from Bruker Biospin, using 0.04ppm width from 0.5 to 10ppm spectral window. The residual water region from 4.68 to 4.88pp was excluded from analysis. All spectra were normalized to the total spectral area and the data table was exported into SIMCA v13.0.3 software for statistical analysis. The PCA analysis was performed using UV-scaling of data and the model was autofit using cross validation rules to determine the number of significant components. The clouds of sample points GF, GM15 and SOPF are distributed according to microbiota complexity from left to right on PC1 (18%) and top to down on PC2 (12%).

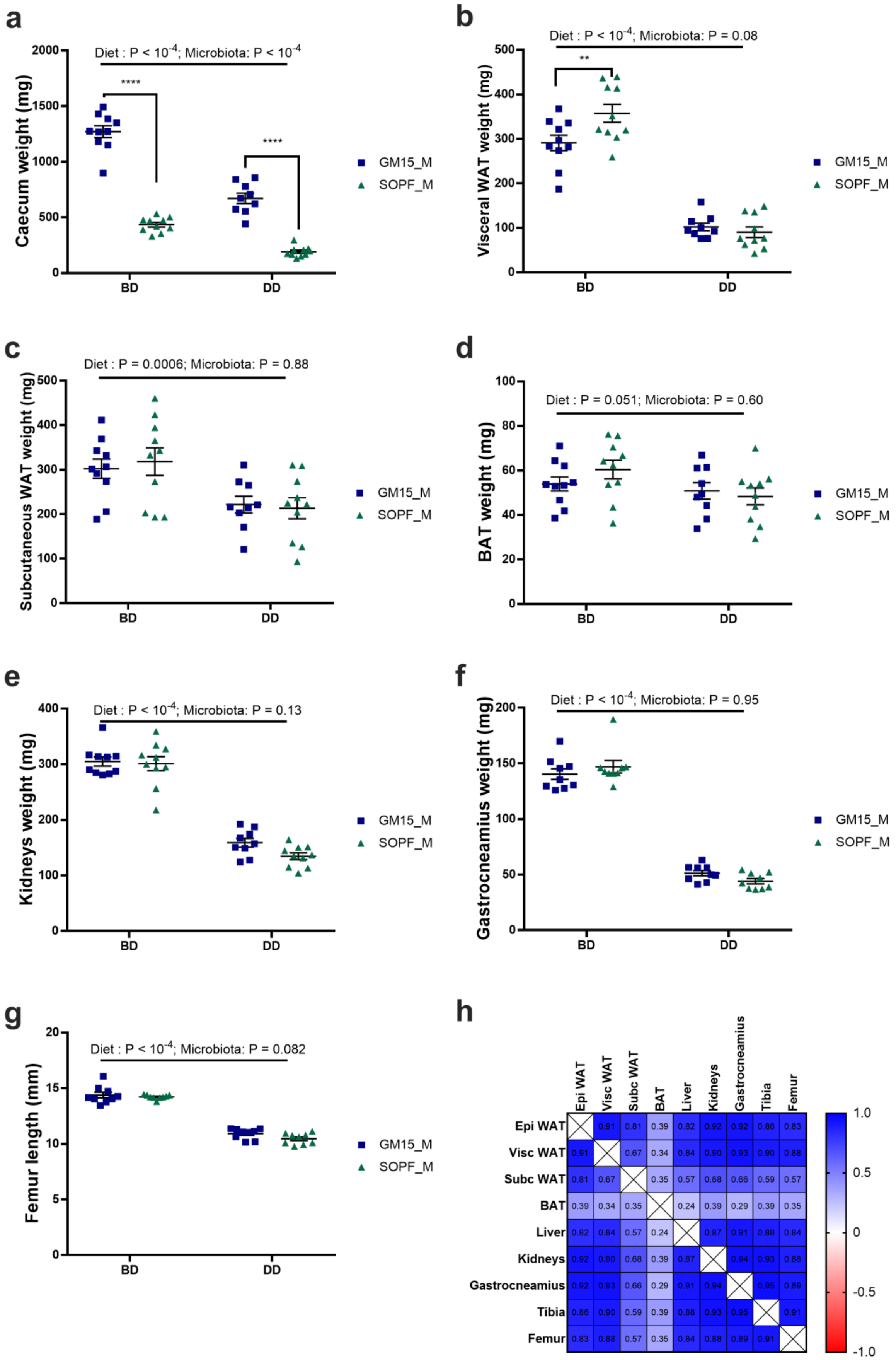
a
Diet : P < 10-4; Microbiota: P < 10-4
****
****
Caecum weight (mg)
2000
1500
1000
500
0
BD
DD
GM15_M
SOPF_M
b
Diet : P < 10-4; Microbiota: P = 0.08
**
Visceral WAT weight (mg)
500
400
300
200
100
0
BD
DD
GM15_M
SOPF_M
c
Diet : P = 0.0006; Microbiota: P = 0.88
Subcutaneous WAT weight (mg)
500
400
300
200
100
0
BD
DD
GM15_M
SOPF_M
d
Diet : P = 0.051; Microbiota: P = 0.60
BAT weight (mg)
100
80
60
40
20
0
BD
DD
GM15_M
SOPF_M
e
Diet : P < 10-4; Microbiota: P = 0.13
Kidneys weight (mg)
400
300
200
100
0
BD
DD
GM15_M
SOPF_M
f
Diet : P < 10-4; Microbiota: P = 0.95
Gastrocneamius weight (mg)
200
150
100
50
0
BD
DD
GM15_M
SOPF_M
g
Diet : P < 10-4; Microbiota: P = 0.082
Femur length (mm)
20
15
10
5
0
BD
DD
GM15_M
SOPF_M
h
Epi WAT
Visc WAT
Subc WAT
BAT
Liver
Kidneys
Gastrocneamius
Tibia
Femur
Epi WAT 0.91 0.81 0.39 0.82 0.92 0.92 0.86 0.83
Visc WAT 0.91 0.67 0.34 0.84 0.90 0.93 0.90 0.88
Subc WAT 0.81 0.67 0.35 0.57 0.68 0.66 0.59 0.57
BAT 0.39 0.34 0.35 0.24 0.39 0.29 0.39 0.35
Liver 0.82 0.84 0.57 0.24 0.87 0.91 0.88 0.84
Kidneys 0.92 0.90 0.68 0.39 0.87 0.94 0.93 0.88
Gastrocneamius 0.92 0.93 0.66 0.29 0.91 0.94 0.95 0.89
Tibia 0.86 0.90 0.59 0.39 0.88 0.93 0.95 0.91
Femur 0.83 0.88 0.57 0.35 0.84 0.88 0.89 0.91
1.0
0.5
0
-0.5
-1.0

**Additional file 9: Fig. S6. Measurements of several organs after nutritional challenge.** Dot plots where dots, lines and error bars represent respectively individual mice (10 males per group), means and SEM. Measurements at day 56 of weights of caecum (**a**), visceral (**b**) and subcutaneous (**c**) white adipose tissue (WAT), brown adipose tissue (BAT) (**d**), kidneys (**e**) and gastrocneamius muscle (**f**) and length of right femur (**g**). P-values after two-way ANOVA were adjusted for Sidak's post-hoc test for multiple comparisons. **h.** Correlation matrix showing Pearson r values of all the measured parameters. For PCA, brown adipose tissue was excluded, given its lack of correlation with other parameters. *P<0.05, **P<0.01, ***P<0.001, ****P<0.0001.

**a**

| GM15 consortium | Facility1_GM15 (N=29) vs. Facility2_GM15 (N=27) F0 and F1 filial generations | Facility1_SOPF (N=19) vs. Facility2_SPF (N=33) 2 consecutive generations |
|---|---|---|
| *Bacteroides acidifaciens* MD185 | ns | ns |
| *Bacteroides caecimuris* MD237 | ** | **** |
| *Parabacteroides goldsteinii* MD072 # | ** | no test (only 2 positive Facility2_SPF samples) |
| *Clostridium cocleatum* I50 * # | ns | ** |
| *Enterocloster clostridioformis* YL32 # | ns | ** |
| *Clostridium* sp. MD294 * # | no test (only 2 positive Facility1_GM15 samples) | ns |
| *Clostridium* sp. MD300 * # | ns (5 positive Facility2_GM15 samples) | ns |
| *Subtilibacillum caecimuris* MD335 ** ## | ns | **** |
| *Longibacillum caecimuris* MD329 ** ## | ns | **** |
| *Irregularicoccus caecimuris* MD308 ** # | ns | ns |
| *Lactobacillus johnsonii* MD006 # | ns | ns |
| *Ligilactobacillus murinus* MD040 | ns | **** |
| *Limosilactobacillus reuteri* MD207 # | ** | no test (only 2 positive Facility2_SPF samples) |
| *Anaerotruncus colihominis* JM4-15 * # | ns (3 positive Facility1_GM15 samples) | ns |
| *Escherichia coli* Mt1B1 | * | * |

**b**

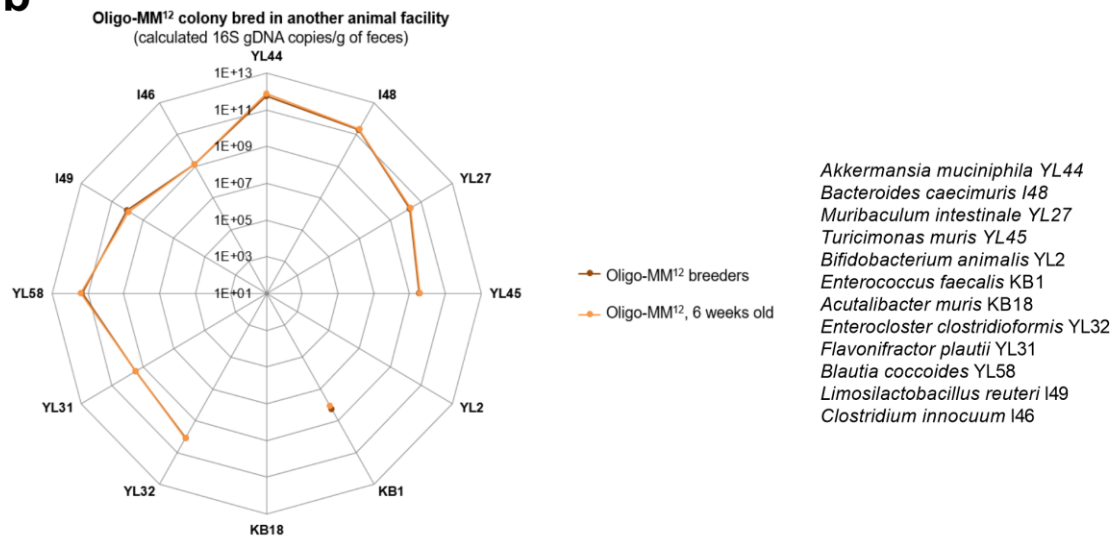


**Additional file 10: Fig. S7. Assessment of gut microbiota reproducibility in GM15 and SOPF/SPF mice in 2 animal facilities, and Oligo-MM$^{12}$ gut microbiota stability in facility 2. a** Summary table of results from Dunn's multiple comparison test applied to GM15 strains concentration quantified by microfluidic qPCR assay in GM15 mice and SOPF/SPF mice between facility 1 and 2. **b** Radar plot showing the Oligo-MM$^{12}$ strains distribution in feces. *Bifidobacterium animalis* YL2 and *Acutalibacter muris* KB18 strains were below qPCR detection limit.

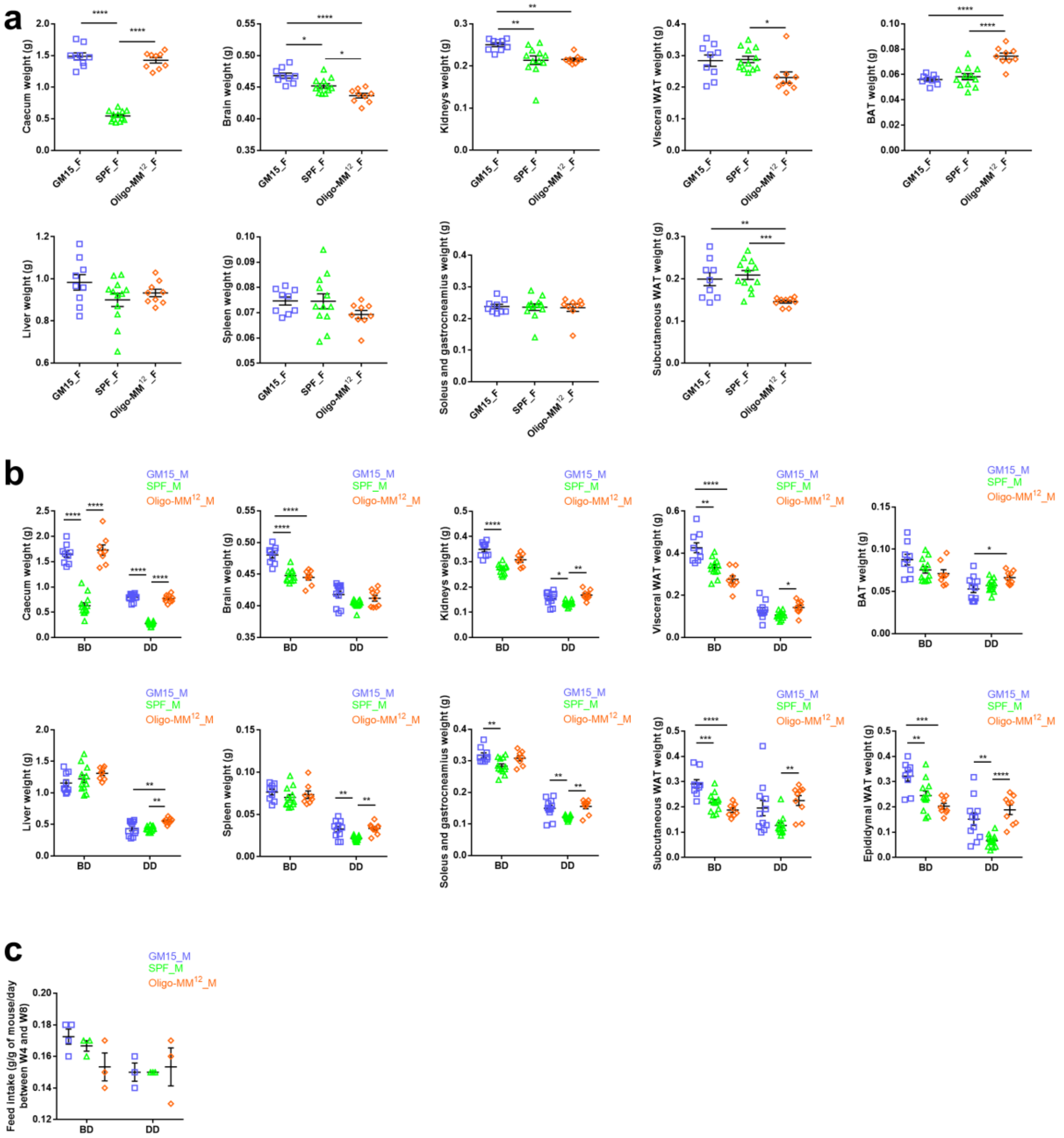


**Additional file 11: Fig. S8. Measurements of organs weight of animals from facility 2. a** and **b** Organs weight. Dot plots where dots, lines and error bars represent respectively individual mice, means and SEM. Female (F) and male (M), breeding diet (BD) and depleted diet (DD), white adipose tissue (WAT) and brown adipose tissue (BAT) respectively. GM15 mice (9 F, 9 M_BD, 11 M_DD), SPF mice (12 F, 12 M_BD, 13 M_DD) and Oligo-MM$^{12}$ mice (9 F, 8 M_BD, 9 M_DD). Dunn's multiple comparison analyses (visceral WAT, kidneys and soleus/gastrocneamius of F; spleen and kidneys of M under BD; brain and subcutaneous WAT of M under DD) or Tukey's multiple comparison analyses otherwise. **c** Feed intake. Dot plot where dots, lines and error bars represent respectively individual cages, means and SEM. Dunn’s multiple comparison analysis. *P<0.05, **P<0.01, ***P<0.001, ****P<0.0001.

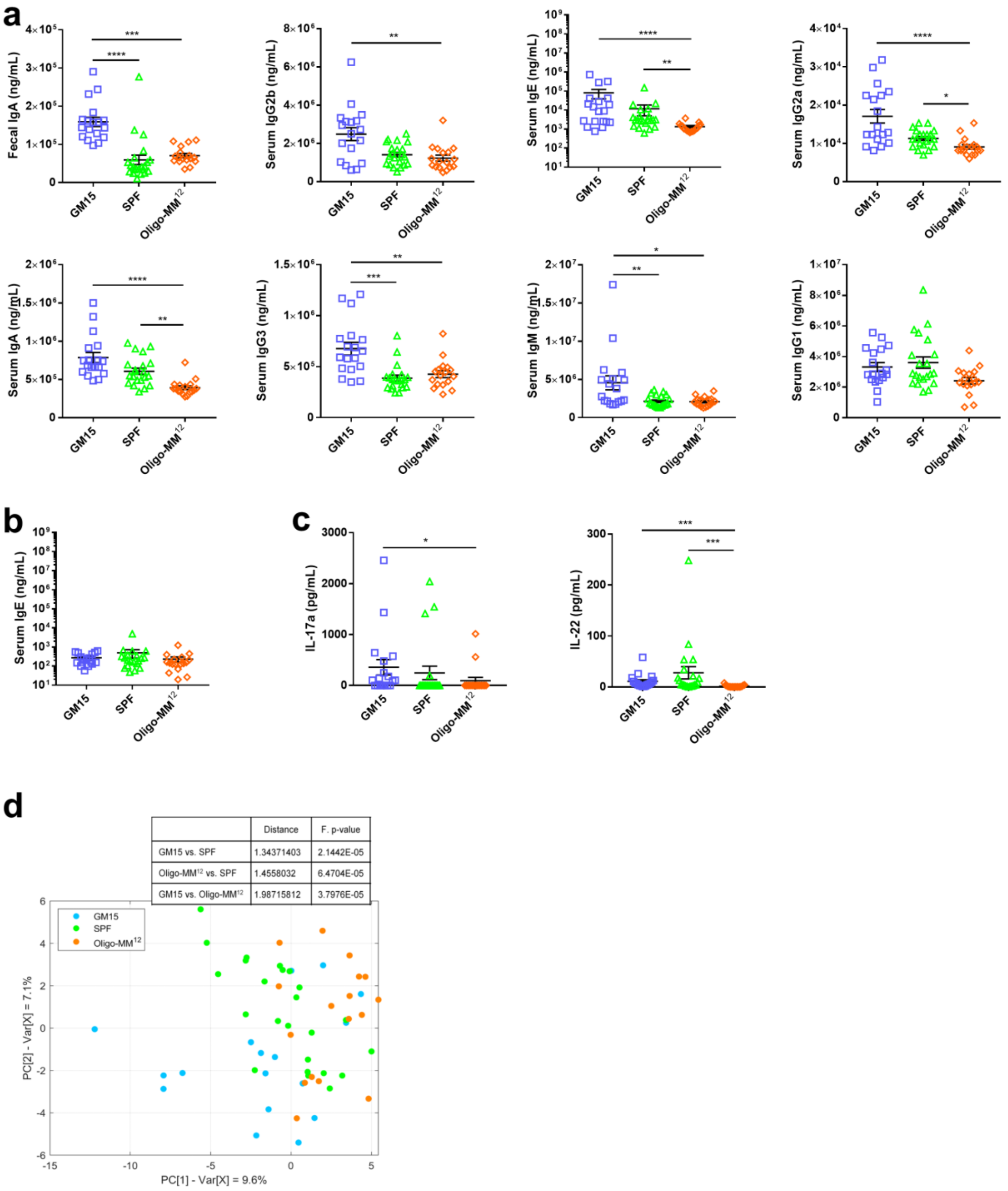


**Additional file 12: Fig. S9. Immune phenotype and metabolic profile of animals from facility 2. a-c** Dot plots where dots, lines and error bars represent respectively individual mice (18 GM15, 24 SPF and 17 Oligo-MM$^{12}$), means and SEM. **a** Fecal IgA, serum IgA, IgG1, IgG2a and b, IgG3, IgE and IgM Luminex analysis. Dunn's multiple comparison analyses. **b** IgE ELISA assay. Dunn's multiple comparison analyses. **c** Circulating IL-17a and IL-22 levels Luminex analysis. Dunn's multiple comparison analyses. **d** PCA score plot representing the distribution of the polar metabolite composition along the two first principal components. *$P<0.05$, **$P<0.01$, ***$P<0.001$, ****$P<0.0001$.
